# AnalyzAIRR: A user-friendly guided workflow for AIRR data analysis

**DOI:** 10.1101/2024.09.25.614939

**Authors:** Vanessa Mhanna, Gabriel Pires, Grégoire Bohl-Viallefond, Karim El Soufi, Nicolas Tchitchek, David Klatzmann, Adrien Six, Hang P. Pham, Encarnita Mariotti-Ferrandiz

**Affiliations:** Sorbonne Université, INSERM, Immunology-Immunopathology-Immunotherapy (i3), F-75005 Paris, France; AP-HP, Hôpital Pitié-Salpêtrière, Clinical Investigation Center for Biotherapies (CIC-BTi) and Immunology-Inflammation-Infectiology and Dermatology Department (3iD), Paris, France; ILTOO Pharma, Statistics Department, F-75013, Paris, France; Institut Universitaire de France (IUF)

**Keywords:** bulk high-throughput sequencing, TCR, BCR, statistical analyses, repertoire diversity, repertoire convergence

## Abstract

The analysis of bulk adaptive immune receptor repertoires (AIRR) enables the understanding of immune responses in both normal and pathological conditions. However, the complexity of AIRR calls for advanced, specialized methods to extract meaningful biological insights. These sophisticated approaches often present challenges for researchers with limited bioinformatics expertise, hindering access to comprehensive immune system analysis. To address this challenge, we developed AnalyzAIRR, an AIRR-compliant R package enabling advanced bulk AIRR sequencing data. The tool integrates state-of-the-art statistical and visualization methods applicable at various levels of granularity. It offers a platform for general data exploration, filtering and manipulation, and in-depth cross-comparisons of AIRR datasets, aimed at answering specific biological questions. We illustrate AnalyzAIRR functionalities using of a published murine dataset of 18 T-cell receptor repertoires from three diferrent T cell subsets. We first detected and removed a major contaminant in a group of samples, before proceeding with to the comparative analysis. Subsequent cross-sample analysis revealed differences in repertoire diversity that aligned with the respective cell phenotypes, and in repertoire convergence among the studied subsets. AnalyzAIRR’s set of analytical metrics is integrated into a Shiny web application and complemented with a tutorial to help users in their analytical strategy, making it user-friendly for biologists with little or no background in bioinformatics.

## Introduction

B-cell receptors (BCR) and T-cell receptors (TCR) are engaged in antigen recognition by B and T cells respectively, sustaining the antigen-specificity of the adaptive immune response. They result from a sophisticated mechanism of genetic recombination of IG (for BCR) or TR (for TCR) genes, forming the adaptive immune receptor repertoire (AIRR). This extremely diverse repertoire carries the immune response capacity of every individual. Thus, AIRRs have been extensively studied to better understand the role of the adaptive immune response in various physiopathological conditions [1, 2, 3], disease progression [4, 5, 6], response to treatment [7, 8] or vaccination [9], [10], and maintenance of homeostasis [11, 12, 13, 14, 15].

High-throughput sequencing (HTS) tools have allowed over the last decade a faster and deeper profiling of AIRRs. Notably, it has permitted the generation of millions of sequences, mostly through bulk sequencing [16, 17, 18], to help characterize the AIRR. This breakthrough led to the advent of sophisticated computational tools [19]. On the one hand, AIRR annotation tools for both TR and IG rearrangement sequencing data were developed, such as MiXCR [20], IMSEQ [21], IgBLAST [22], and pRESTO from the Immcantation suite [23]. These tools allow the mapping of V, D, and J genes, the extraction of the complementary determining region 3 (CDR3), and the assembly and counting of AIRR rearrangements, namely clonotypes. On the other hand, strategies have been adapted from various fields such as ecology and information theory to describe, quantitatively and qualitatively, different aspects of the AIRR [24, 25]. Such strategies, including diversity estimation and gene usage calculation, are implemented in a list of tools such as ImmuneREF [26], Immunarch [27], and Immcantation [28].

Despite the wide range of analytical strategies proposed by most of these tools, they all require bioinformatics and programming skills, limiting the accessibility for non-bioinformatician researchers. Therefore, there is a need to better inform the scientific community of how and when to use the different suitable metrics for accurate AIRR studies.

Herein, we present AnalyzAIRR, an R package that proposes a systematic analytical routine to guide bioinformatics expert and non-expert users in AIRR analytical strategies. The package offers various statistical metrics and visualization methods, allowing a complete data exploration as well as cross-sample comparisons to answer defined biological questions. Advanced users can customize the visualizations generated by AnalyzAIRR in terms of style to better align with their preferences. Additionally, they can extract statistical data for use in other visualization platforms or tailor the presentation of the data to address specific research needs. Importantly, we complemented AnalyzAIRR with an intuitive R-Shiny web framework, making it accessible to biologists with limited bioinformatics expertise. This user-friendly interface allows them to explore and implement the proposed analytical strategies, bridging the gap between complex data analysis and practical research applications.

## Design and implementation

### Functionalities

AnalyzAIRR integrates a comprehensive set of methods for IG/TR repertoire analysis into an R package, accessible to both bioinformatics experts and non-experts. Three key functionalities are offered by AnalyzAIRR.

### Comprehensive data exploration

AnalyzAIRR provides tools for the examination of individual samples or the entire dataset. This is achieved through the calculation of a set descriptive satistics including counts of sequences, genes, and clones, and measures of clonal distribution and diversity. This process facilitates the detection of potential outliers or contaminants within the data.

### Advanced data manipulation

This functionality allows to filter out specific sequences or samples based on user-defined criteria. In addition, two normalization methods are proposed to ensure comparability across samples: i) random down-sampling, and ii) exclusion of low-count sequences using Shannon entropy [29]. The latter further allows the correction of the clonotype distribution in small samples which are often over sequenced.

### In-depth cross sample comparisons

AIRR comparisons between sample groups are possible through staregies such as, diversity estimation, repertoire similarity evaluation and the identification of differentially expressed genes/clonotypes between experimental groups. These comparative analyses enable the identification of significant patterns and differences, helping researchers address specific biological questions.

### Interactive AIRR data analysis interface

AnalyzAIRR’s set of analytical metrics is integrated into a Shiny web application, which makes it user-friendly for biologists (**Error! Reference source not found.**). Users are can apply the full range of analytical methods proposed by AnalyzAIRR, without requiring any bioinformatics expertise. They are guided by a detailed documentation that explains each function and provides relevant use cases.

The Shiny includes additional features, such as interactive plots that reveal information that might be missed in static visualizations. Furthermore, users can generate a summary report compiling all plots created during an analysis session. This report can be customized with descriptive text for each figure, allowing for detailed interpretation of results, and can be downloaded in various formats, including PDF, HTML, and PowerPoint.

### Data input and structure

AIRR sequencing datasets include a collection of V(D)J rearrangements, either IG or TR, also named clonotypes. Their analysis requires primarily an annotation step using dedicated alignment tools to transform raw sequencing data into analyzable immune repertoire information. AnalyzAIRR supports input files: (i) generated by a list of annotation tools including MiXCR and immunoSEQ; (ii) formatted following the MiAIRR standard guidelines from the AIRR Community [30]; or (iii) formatted following AnalyzAIRR requirements. A descriptive metadata providing sample information, such as experimental group, age, genetic background, sample type, and cell subset, can also be supplied allowing intra- and inter-group analyses.

AnalyzAIRR employs an R-structured data container, known as an S4 object, that ensures efficient manipulation of the complex AIRR-seq data throughout the analysis workflow. AnalyzAIRR’s object, named RepSeqExperiment, can be created using the annotated AIRR data files and the metadata, when available, and is used as data input for all AnalyzAIRR functions. Alternatively, users can upload annotated files and metadata directly into the Shiny interface, which will automatically create and load the RepSeqExperiment object to be used in their analysis.

The RepSeqExperiment object is composed of four compartments, each designed to hold a specific type of data: (i) assayData, which encloses all the clonotype tables in the dataset; (ii) metaData, which contains the sample information provided by the user as well as a set of statistics calculated for each sample including the number of sequences and clones; (iii) otherData, which contains the output of many filtering and calculation functions further exemplified in the next section; and (iv) History, which registers all the actions performed on the RepSeqExperiment object.

## Results

To illustrate the above-mentioned AnalyzAIRR functionalities, we made use of a published dataset composed of 18 TCR repertoires of effector T cells (Teff), naïve regulatory T cells (nTreg), and activated/memory regulatory T cells (amTreg) collected from the spleen of young C57BL/6 mice [8]. The files were aligned with MiXCR (v.4.4.1) and used as input along with the corresponding metadata file. Herein, we will analyze the TRA repertoire of the 18 samples, as a RepSeqExperiment object can be generated for one chain at a time.

### Data validation process

AnalyzAIRR proposes a series of analyses that can be performed on each sample within the dataset, enabling an exploratory description of the dataset. Available metrics include descriptive statistics such as the number of sequences and the number of clones (corresponding to each unique V-CDR3aa-J combination), the Renyi entropy, which assesses various repertoire diversity metrics [31], the clonal distribution and the V and J gene usage. These approaches can serve first as a tool to perform dataset validation, an essential step in processing AIRR-seq data.

To assess the clonotype richness and the sequencing depth of a sample, the Chao1 index and rarefaction curves serve complementary roles. Rarefaction curves assess the sufficiency of sequencing depth and sampling (**Fig 1A**); A plateau in the curve suggests that sampling is yielding few or no new species, whereas a steep ascending curve indicates that additional sequencing may still yield new clones. In our example data, sequencing depth was sufficient in samples that show a flatenned curve, while further sequencing could be required for ones showing a steep curve.

**Fig 1.**
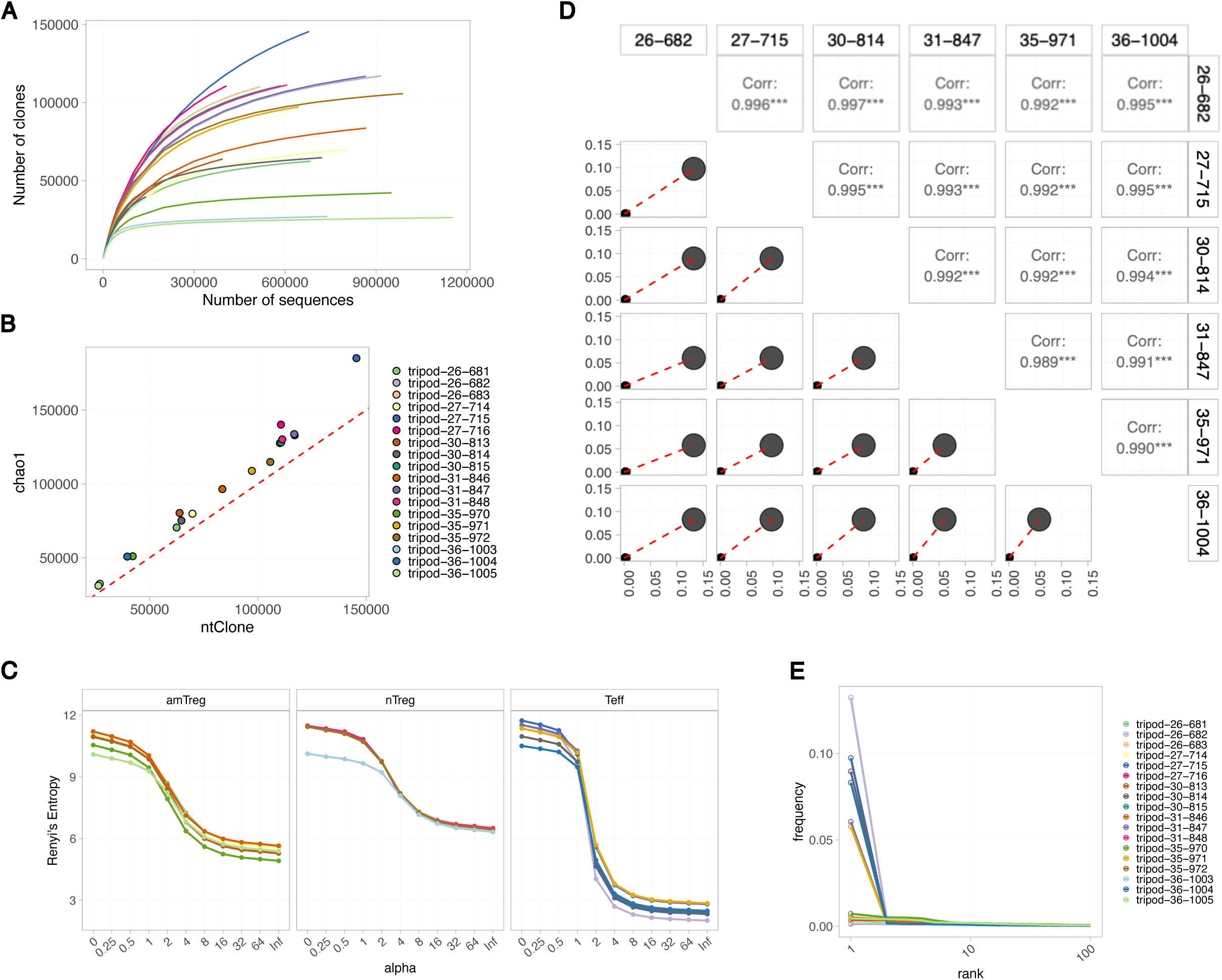
Exploratory analysis of the full dataset. A) Rarefaction curves are plotted for each sample, illustrating the relationship between the number of sequences randomly selected from the sample and the number of clones they represent. Each sample is represented by a unique color. B) The Chao1 richness estimation is plotted as a function of the observed number of ntClones in each sample. The diagonal dashed line represents the reference line. C) Renyi values were calculated at the aaClone level for each of the pre-defined alpha values and plotted for each sample. Samples are separated based on the cell subset to which they belong. D) Scatter diagrams are generated for every Teff sample pair. Lower parts show scatter points representing the aaClones and their corresponding frequencies in each compared sample. Each circle corresponds to a unique aaClone. Upper parts show the Pearson correlation calculated on the symmetrical scatter plot, and the significance stars: ***: p-value < 0.001; **: p-value < 0.01; *: p-value < 0.05 and.: p-value < 0.10. E) Curves plotting the aaClone frequencies as a function of their occurrence rank within each sample. Only the top 100 clones are plotted.

To calculate the number of clones that are not observed in the sample, and that could be detected with further sampling and/or sequencing, the Chao1 index can be used. This index estimates the total clone richness of a sample including clones that are likely present but unseen or undetected (**Fig 1B**). Consistently, samples with a high Chao estimation were ones showing a steep rarefaction curve. Together, both metrics provide a comprehensive understanding of species richness and sampling adequacy in the different samples.

Furthermore, assessing the repertoire diversity of the observed clones using the Renyi curves allowed us to detect a contaminant clone, present in high occurrences in the Teff samples. Indeed, the Renyi curves showed a sharp decline in a group of samples, all being Teff cells (**Fig 1C**). This observation is not expected in such a polyclonal and diverse cell subset. In addition, nearly perfect clone frequency correlations were observed between each pair of Teff samples, caused by one shared clone present at a significant higher frequency than the other clones (**Fig 1D** and **E**). This is once more an unusual observation in polyclonal Teff repertoires collected from different mice.

### Single sample analysis and filtering

In view of these observations, and to confirm the validaty of our results, we examined each of the problematic Teff sample individually. The analysis performed on one Teff sample revealed a decrease in its repertoire diversity, as evidenced by both the Simpson and Berger-Parker indices, compared to the other samples within the dataset (**Fig 2A**). This reduced diversity can be explained by the high cumulative frequency of clones with occurrences higher than 10,000 (**Fig 2B**), suggesting significant clonal expansions. Specifically, these expansions were driven by a single dominant clone, which exhibited a significantly higher count than subsequent clones within the same repertoire (**Fig 2C**). This clone expressed one V-J combination, thus altering the overall distribution of V and J genes within the sample (**Fig 2D**). These along with the previous results allowed the identification of the contaminant clone sequence, which was filtered out from all Teff samples. The filtering rescued the diversity and the clonal sharing in all Teff samples (**Error! Reference source not found.**).

**Fig 2.**
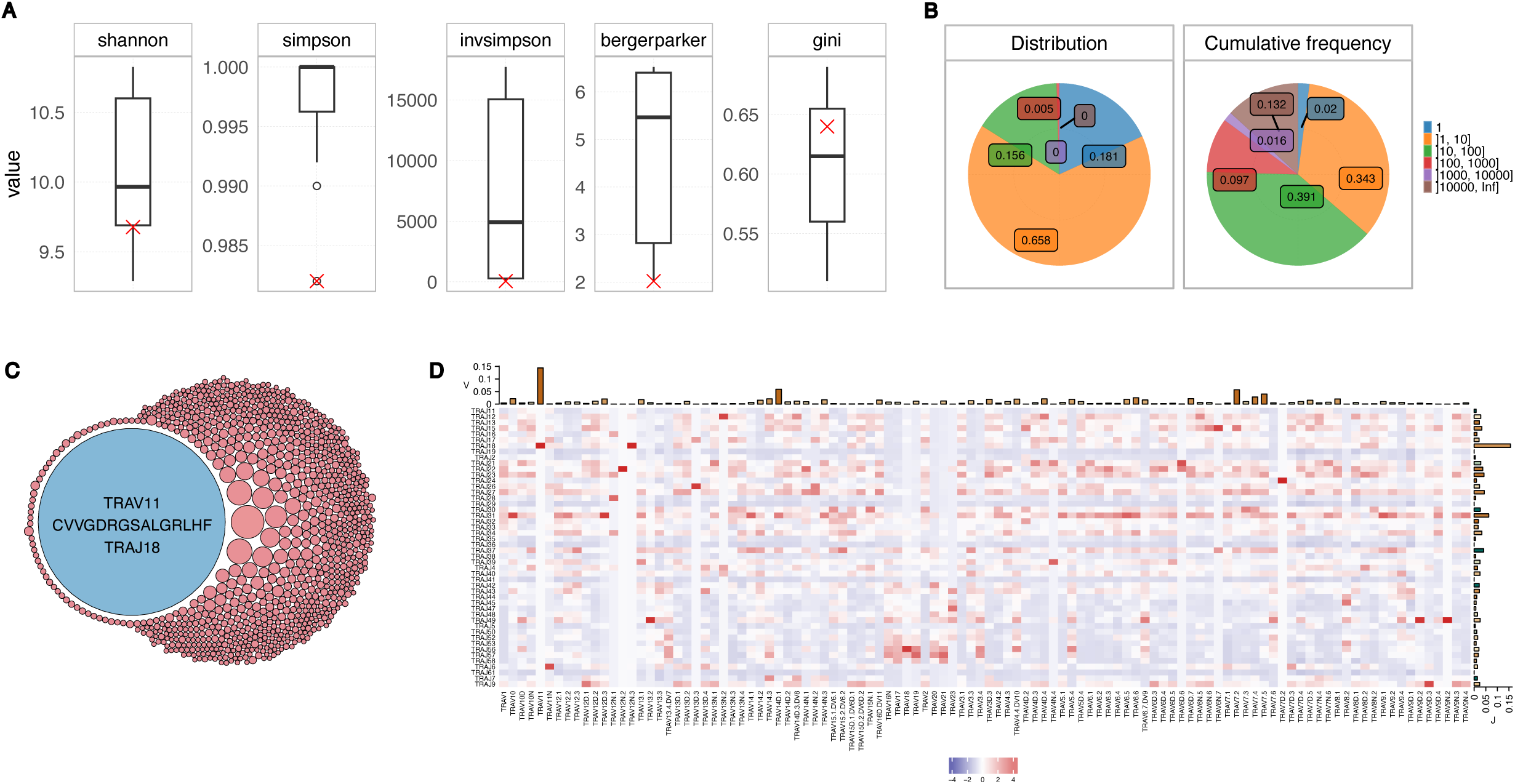
Single Teff sample analysis. A) Boxplots showing different diversity indices calculated on all the dataset samples. The sample of interest “tripod-26-682” is represented by a red cross. B) Pie charts representing the distribution (left) and the cumulative frequency (right) of clones within each count interval for the sample of interest. C) A circular treemap plotting the top 1% aaClones in the sample of interest. Each circle corresponds to a unique clone, and its size is proportional to the clone count. This plot shows the clone distribution within the sample repertoire. D) Heatmap of V and J combination usage within the sample of interest. Frequencies, represented by the color scale, are scaled column-wise. Barplots at the top and right side of the heatmap show the usage of each gene across the row or the column for the V and J genes, respectively.

### Sample normalization

Comparing AIRR-seq data collected from different sources can often result in repertoires of differing sizes, due to biological or technical factors. A key feature of AnalyzAIRR is its ability to normalize data, ensuring comparable sample sizes and accurate comparisons in AIRR-seq datasets. AnalyzAIRR offers, among others, a Shannon-based down sampling method, leveraging the Shannon entropy to exclude sequences considered as noise [29]. While this normalization process resulted in a decrease in repertoire richness and diversity across all our samples (**Fig 3A** and **B**), the pre- and post-normalization repertoires clustered per sample based on the Jaccard scores, which measures the similarity between two samples independently of the species abundance [32] (**Fig 3C**). With the normalization applied; the datasets are now prepared for extensive comparative analysis.

**Fig 3.**
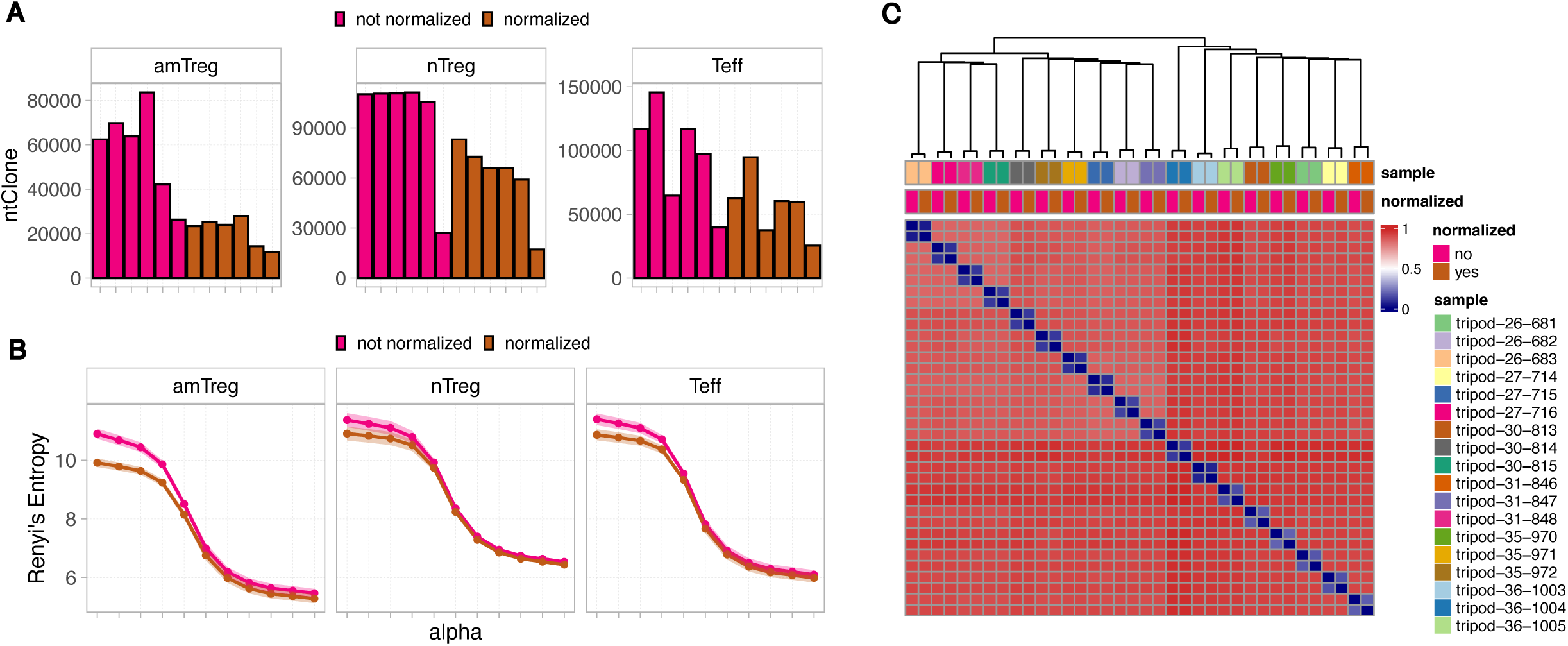
The effect of the Shannon normalization on the repertoires. A) Barplots showing the number of nt clones (V-CDR3nt-J combinations) for each of the not normalized (pink) and normalized (brown) samples. B) The Renyi values were calculated at the nt clone level at pre-defined alpha. For each of the not non-normalized and normalized groups, the mean (circles) and standard error (shade) are plotted. C) A heatmap showing the Jaccard distances calculated at the aa clone level between all pairwise samples. Samples are labeled column-wise based on the sample name and the normalization condition. Scores range from 0 (complete similarity) to 1 (complete dissimilarity). The Ward’s method was used to perform the hierarchical clustering.

### Gene Usage Analysis

Users can leverage AnalyzAIRR to investigate biological hypotheses, by drawing a detailed comparative analysis between experimental groups and conditions. The first layer of any AIRR-seq data analysis involves comparing the gene usage between conditions. In our dataset, for visual clarity, we compared TRAV and TRAJ usages between the naïve and activated Treg cells. Comparing the gene frequencies showed significant differences in the usage of many gene families (**Fig 4A** and **B**). Additionally, a differential expression analysis performed at the V-J combination level enabled the identification of differentially expressed gene combinations, contributing to the separation of the two subsets in the MDS analysis (**Fig 4C** and **D**).

**Fig 4.**
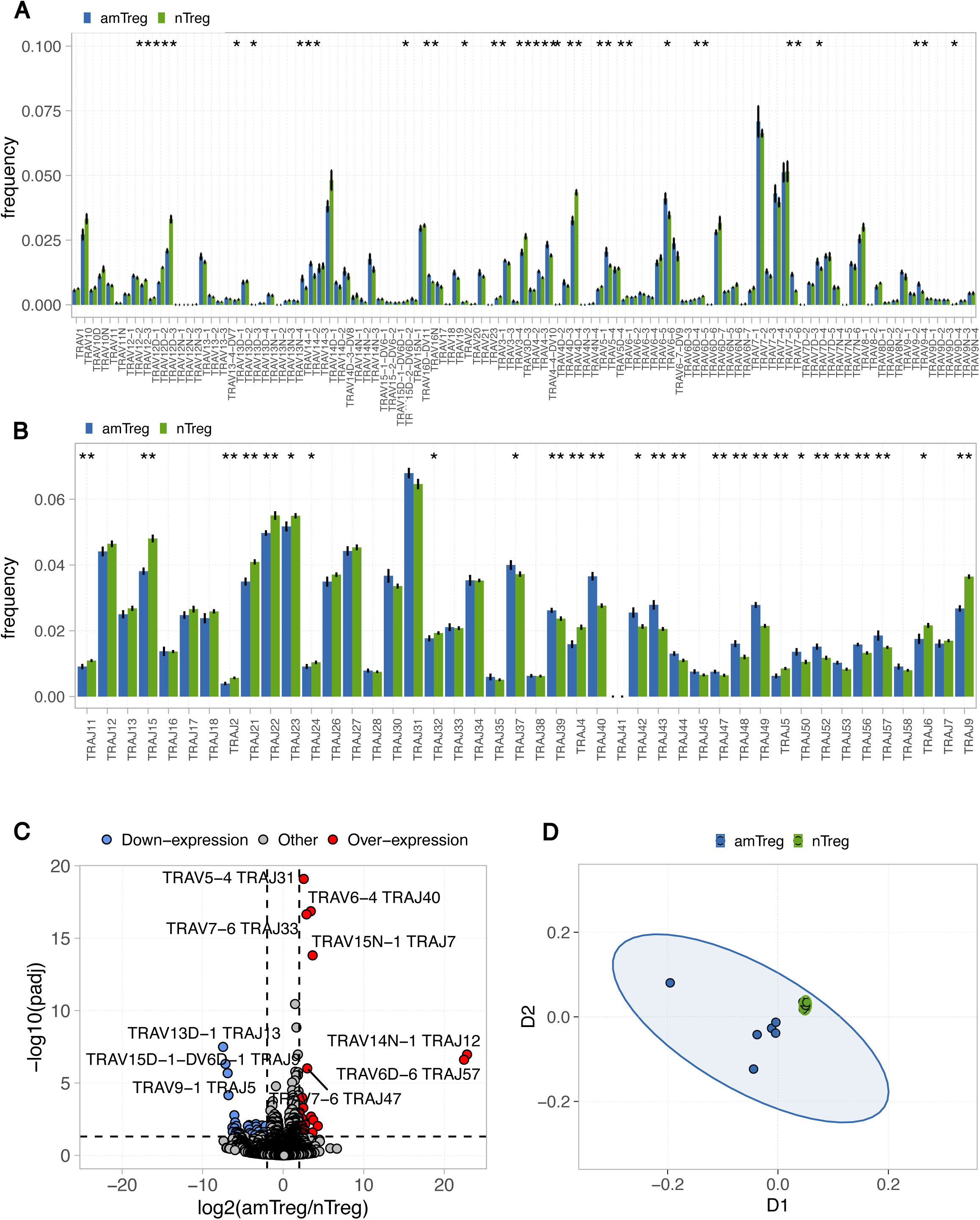
Gene usage analysis between the Treg subsets. Comparison of the TRAV (A) and TRAJ (B) frequencies between amTregs (blue) and nTregs (green) samples. Lines on barplots correspond to the standard error values. A Wilcoxon test was performed and only significant values, when present, are plotted. C) A volcano plot showing differentially expressed TRAV-TRAJ combinations between amTreg and nTreg samples. Over-expressed combinations in the amTreg population are shown in red, down-expressed one in blue based on the fixed thresholds: a fold change > +2 and a pvalue ≤ 0.05. Only the 10 most differentially expressed combinations are labeled. D) Morisita-Horn distances was calculated on TRAV-TRAJ counts between all pairwise Treg samples and used to perform a multidimensional scaling analysis on a two-dimensional space. Each point corresponds to a sample and is colored based on the cell subset it belongs to; amTregs in blue and nTregs in green.

### Clonality and diversity assessment between conditions

We next compared the repertoire clonality of the three different cell subsets. Results showed higher clonality in Teffs and nTregs compared to amTregs, with a significant difference between amTregs and Teffs (**Fig 5A**). We also examined any potential sex-related differences, revealing comparable clone numbers in male and female mice in each cell population (**Fig 5B**). Furthermore, Renyi curves showed distinct profiles, with nTregs exhibiting the most diverse repertoire and amTregs the least diverse (**Fig 5C**), in agreement with the clonality results. Finally, we confirmed this observation by computing and comparing the cumulative frequency of expanded clones, which were found to be higher in amTregs compared to the other subsets (**Fig 5D**).

**Fig 5.**
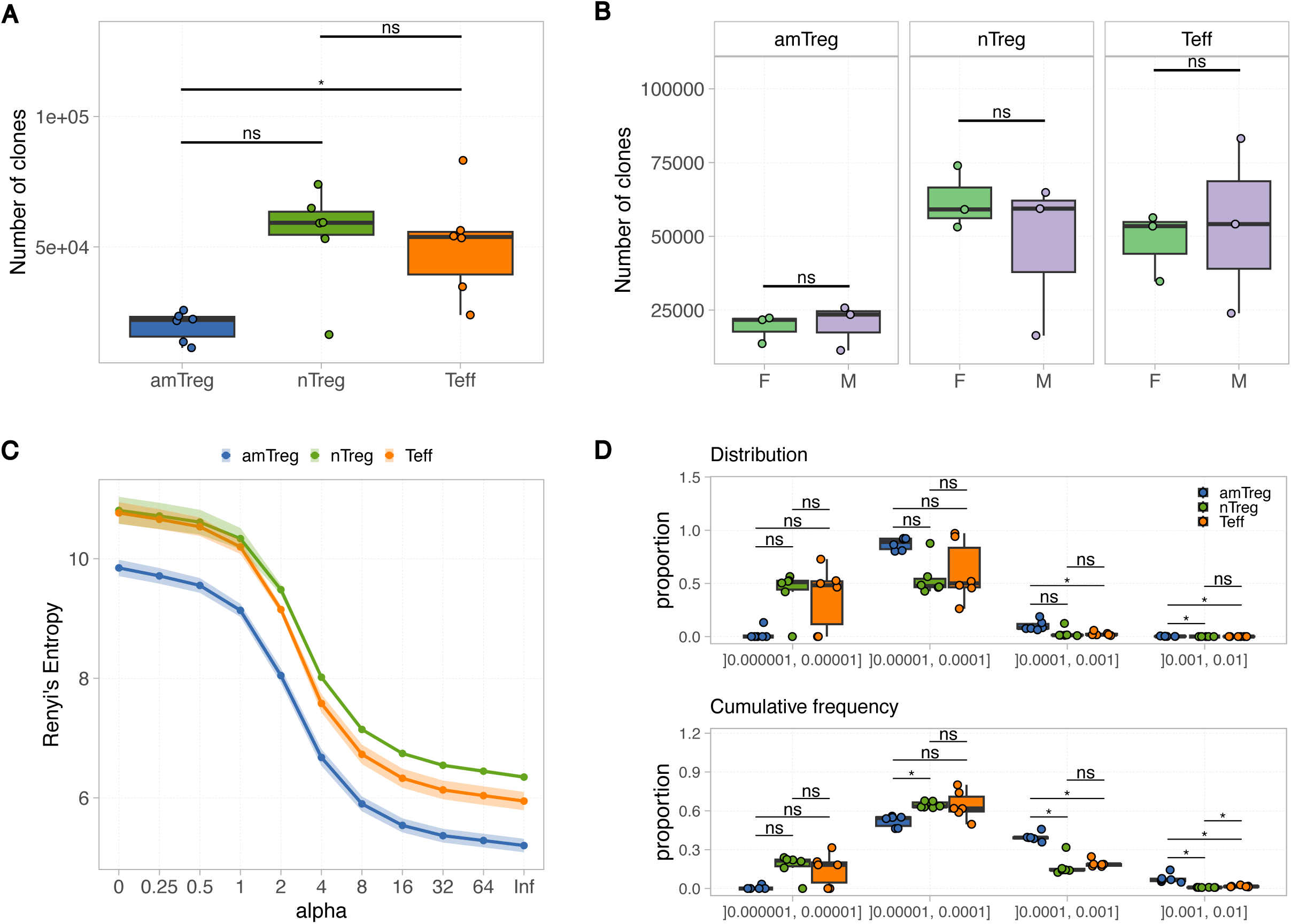
Comparison of the repertoire diversity between the three cell subsets. A) The number of aaClones (bottom) is compared between amTreg (blue), nTreg (green) and Teff (orange) samples. B) The number of aaClones is compared between samples collected from female (green) and male (purple) mice within each cell subset. A-B) Boxplots represent the median across all samples belonging to the same group. A Wilcoxon test is applied and adjusted p-values using the Holm method are shown. C) The Renyi values were calculated at the aaClone level at the pre-defined alpha values. For each specified group, the mean (circles) and standard error (shade) are plotted. D) The distribution (top) and the cumulative frequency (bottom) of clones within each clone fraction interval was compared between the three cell subsets. Boxplots represent the median across all samples belonging to the same group. A Wilcoxon test is applied and adjusted p-values using the Holm method are shown. *P<0.05; **P<0.01; ***P<0.001; ****P<0.0001; ns P>0.05.

### Repertoire composition comparison

The similarity metrics proposed by AnalyzAIRR enabled us to examine the repertoire composition between the different cell subsets, providing insights into repertoire convergence. For instance, hierarchical clustering based on the Jaccard distance separated amTreg samples from Teff and nTreg samples, despite the high dissimilarity scores that indicate modest differences in repertoire compositions (**Fig 6A**). Multidimensional scaling (MDS) analysis applied to the calculated Jaccard scores, also facilitated by AnalyzAIRR, confirmed these findings (**Fig 6B**). In contrast, the Morisita-Horn index, which considers and is sensitive to species abundance, allowed clear discrimination between the three cell subsets in both hierarchical clustering and MDS analysis (**Fig 6C** and **D**). Moreover, calculating the inter-sample similarity scores revealed that nTregs had the most public repertoires, while amTregs had the most private ones.

**Fig 6.**
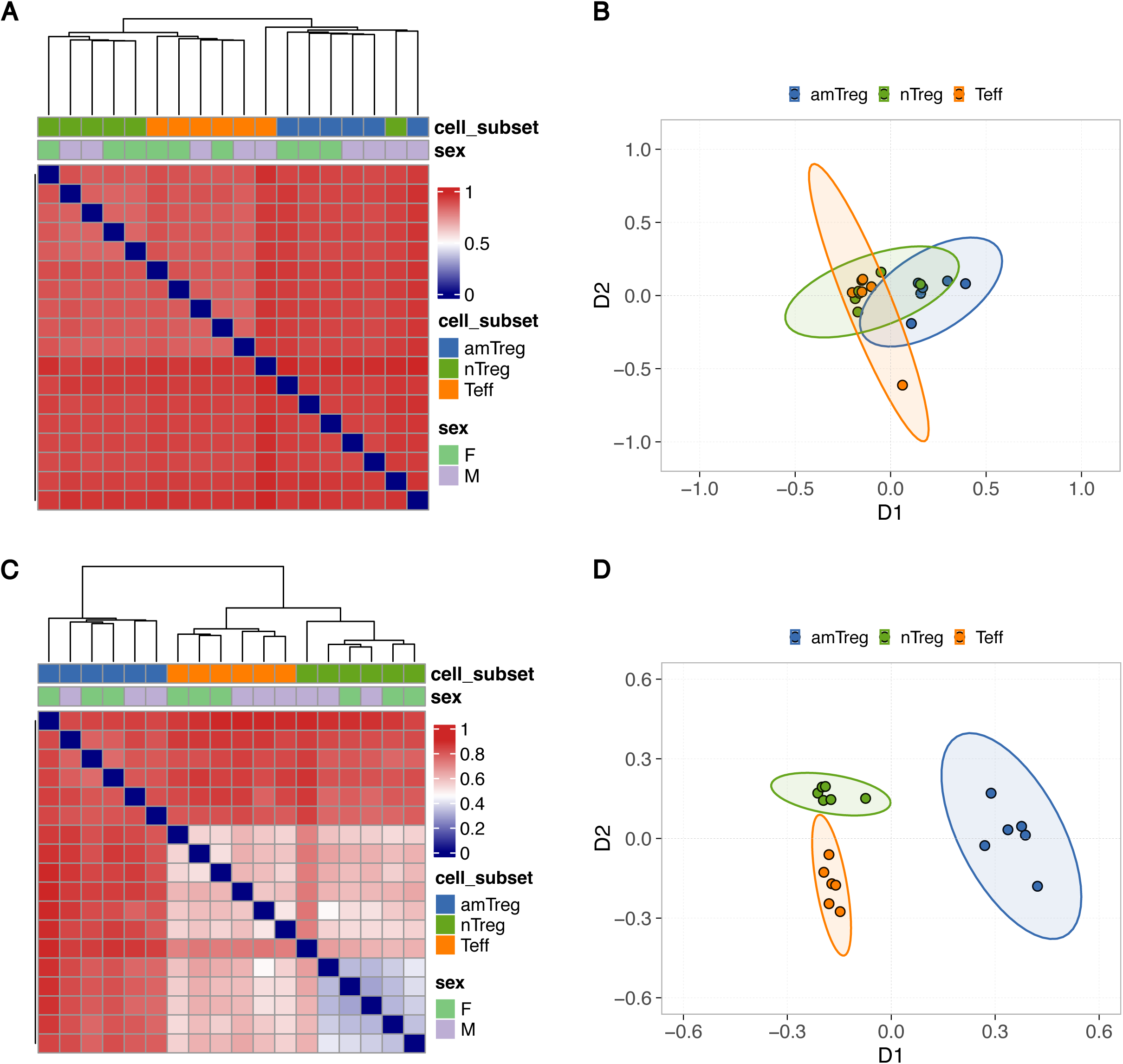
Comparison of the repertoire composition. A heatmap showing the Jaccard (A) and Morisita-Horn (C) distances calculated at the aaClone level between all pairwise samples. Scores range from 0 (complete similarity) to 1 (complete dissimilarity). Samples are labeled column-wise based on the cell subset and sex of the mice used in the study. The Ward’s method was used to perform the hierarchical clustering. The calculated distances were used to perform a multidimensional scaling analysis on a two-dimensional space (B and D). Each point corresponds to a sample and is colored based on the cell subset it belongs to; amTregs in blue, nTregs in green, and Teffs in orange. Ellipses are drawn around points belonging the same group.

## Conclusion

AnalyzAIRR equips researchers with a scalable, AIRR-compliant framework accompanied with a detailed documentation for in-depth exploration of immune receptor repertoires. Using the source code or the Shiny web interface, users can navigate a sequential analysis pipeline to ensure dataset quality, explore individual samples, and perform cross-sample comparisons. In this study, we introduced the various analyses offered by AnalyzAIRR through a direct application on a published TCR-seq dataset. By leveraging its functionalities, we were able to illustrate relevant biological insights, validating the package’s capability to support both expert and non-expert users in the field in conducting comprehensive repertoire analyses. Compared to existing tools, AnalyzAIRR distinguishes itself by offering comprehensive analytical features coupled with a user-friendly interface (**Fig 7**). This combination makes it an ideal tool for both experts and novices in the AIRR-seq or the bioinformatics field.

**Fig 7.**
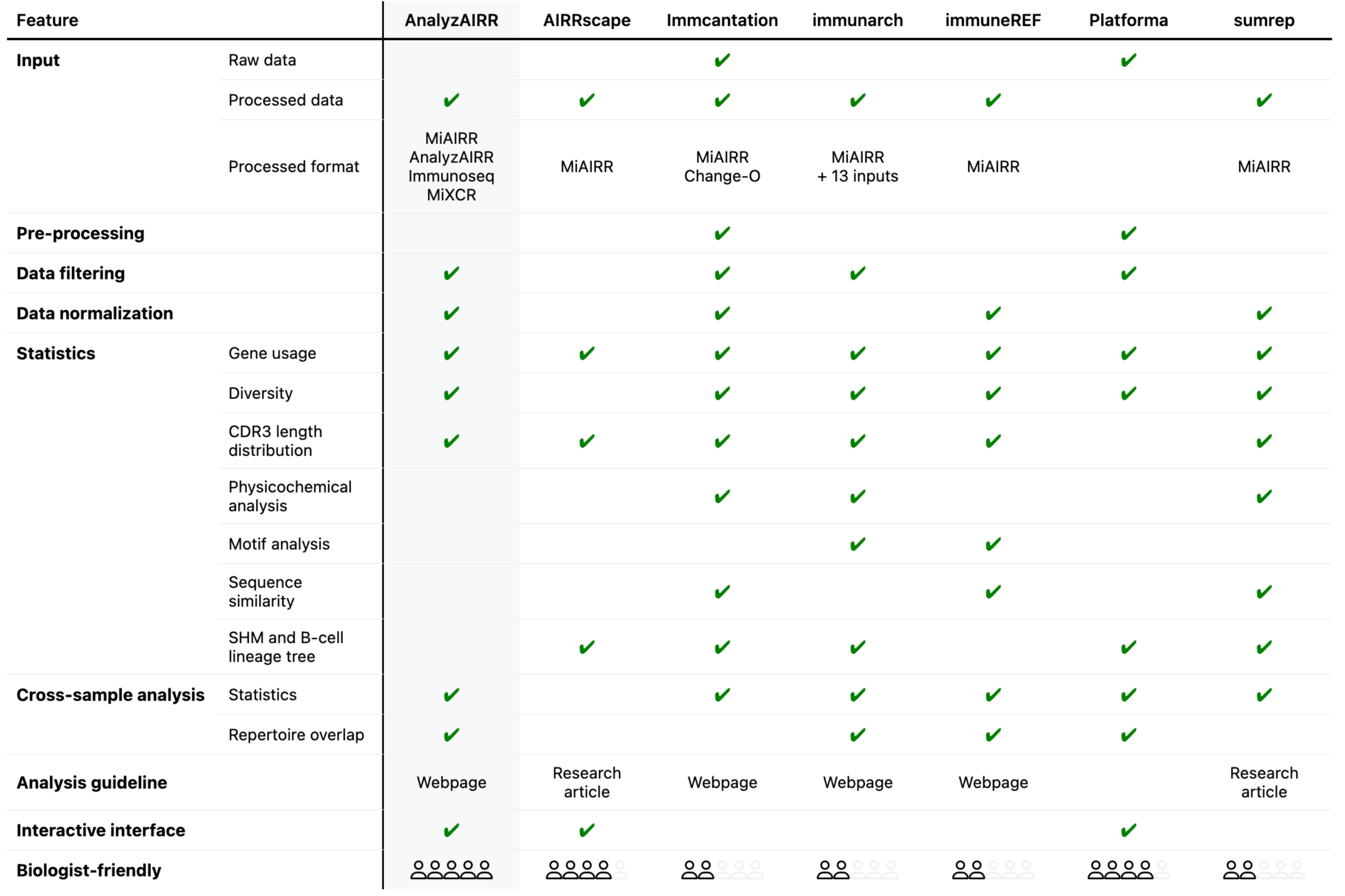
Overview of community-developed tools for the analysis of bulk AIRR-seq data. This table provide a comparative overview of the most used community-developed tools for analyzing bulk AIRR-seq data. The list of criteria includes input data type and pre-processing, data filtering and manipulation, exploratory and cross-sample analysis, somatic hypermutation and phylogenetic (lineage tree) analysis, the availability of a tool documentation, presence of an interactive interface and easy-of-use score for biologists.

## Bibliographical note

Dr. Vanessa Mhanna, PhD, is a postdoctoral fellow at Sorbonne Université, specializing in systems immunology.

Gabriel Pires, MSc, is a bioinformatician at Discngine, specializing in OMICS analytics solutions.

Grégoire Bohl-Viallefond, MSc, is a PhD student at the Gregor Mendel Institute of Molecular Plant Biology, Vienna, Austria, specializing in epigenetics and bioinformatics.

Dr. Karim El Soufi, PhD, is a lead cloud architect at JCDecaux.

Dr. Nicolas Tchitchek, PhD, is an associate professor of systems immunology at Sorbonne Université.

Prof. David Klatzmann, MD-PHD, is a professor of immunology at Sorbonne Université. Prof. Adrien Six, PhD, is a professor of systems immunology at Sorbonne Université.

Dr. Hang-Phuong PHAM, PhD, is a chief data science officer at Parean Biotechnologies, specialized in computational biology.

Prof. Encarnita Mariotti-Ferrandiz, PhD, is a professor of systems immunology at Sorbonne Université.

## Key points

- AnalyzAIRR integrates state-of-the-art statistical and visualization methods for in-depth exploration of bulk AIRR-seq data.
- AnalyzAIRR’s key functionalities include general data exploration, advanced data manipulation and in-depth cross-comparisons of AIRR datasets.
- AnalyzAIRR provides a guided analytical workflow and an interactive interface for biologists with limited bioinformatic skills.
- AnalyzAIRR bridges the gap between complex data analysis and practical research applications.

## Availability

AnalyzAIRR is implemented in R version 4.2.2 (License: GPL-3) and available for installation at: https://github.com/i3-unit/AnalyzAIRR. A vignette providing installation details, function documentation and use cases is also available at: https://github.com/i3-unit/AnalyzAIRR.github.io. The shiny version of the tool is available at https://i3-unit.github.io/AnalyzAIRR/. The example data analysed in this paper and the code developed to generate the plots are publicly available at Zenodo under the DOI: 10.5281/zenodo.13350295.

## Acknowledgments

We thank Pierre Barennes, Valentin Quiniou and Wahiba Chaara for their contribution in conceiving the package.

## Fundings

This work was funded by the TRiPoD European Research Council-Advanced EU (No. 322856) and iMAP (ANR-16-RHUS-0001) grants to DK, the iReceptorPlus (Horizon 2020 No. 825821), the AIR-MI (ANR-18-ECVD-0001) grants to EMF with additional support from the Sorbonne Université, INSERM and AP-HP. ILTOO Pharma supported HPP.

## Declaration of interest

There is no duality of interest relevant to this article.

## Notes

### Competing Interest Statement

The authors have declared no competing interest.

