## Supplemental Figures for "AnalyzAIRR: A user-friendly guided workflow for AIRR data analysis"

**AnalyzAIRR: A user-friendly guided workflow for AIRR data analysis**
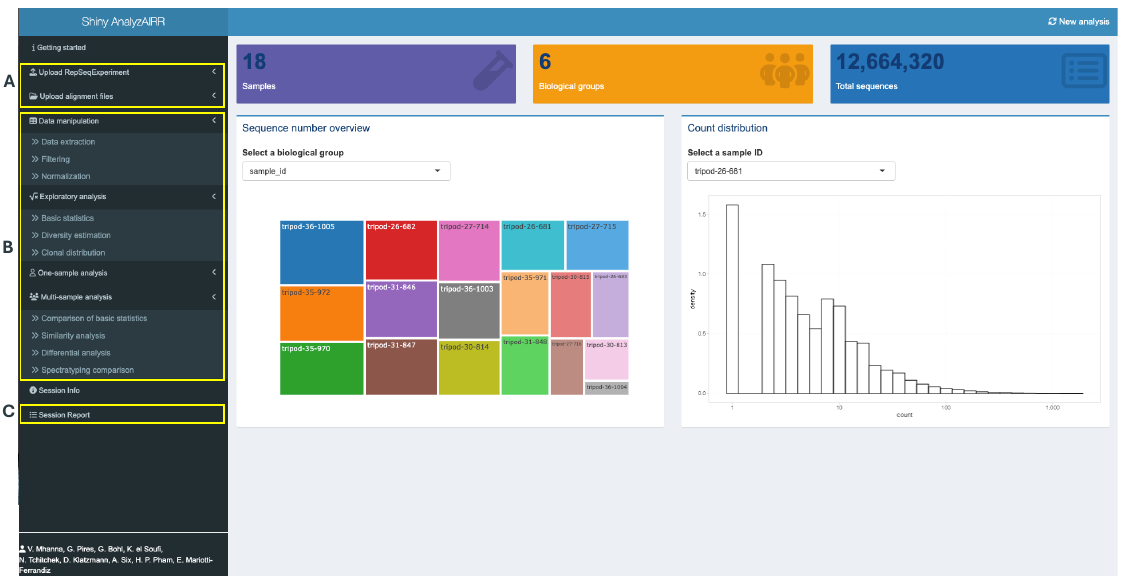

**S1 Fig.** **The Shiny-AnalyzAIRR interface**. A) Aligned file formats supported by AnalyzAIRR can be loaded directly into the interface, which will create a RepSeqExperiment object to be used in the analysis. The interface also allows the upload of a source-code-generated RepSeqExperiment object. B) The four main features of AnalyzAIRR include a data exploration step, data manipulation, one-sample exploration, and cross-sample analysis. C) All generated plots in an analysis session are compiled in a summary report that can be commented on and downloaded.

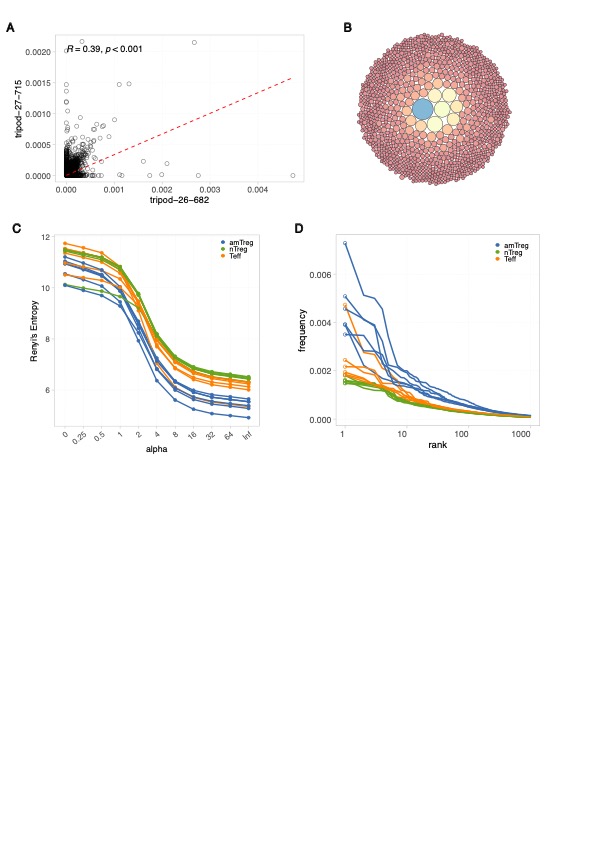

**S2 Fig.** **Filtering out the identified contaminant clone.** A) Scatter plot of aaClone frequencies between two Teff samples post-filtering. A linear regression model is fitted onto the data and represented by the red dashed line. The Pearson correlation is plotted. B) A circular treemap was generated for sample “tripod-26-682” and showed the structure of the top 1% of aaClones in the sample repertoire. Each circle represents a unique aaClone, and the circle size corresponds to the clone count. C) The Renyi values were calculated at the aaClone level for each of the pre-defined alpha values and plotted for each filtered sample. D) The distribution of clones as a function of their occurrence rank within each sample after the removal of the contaminant clone. amTreg samples are represented in blue, nTregs in green, and Teffs in orange.
